# DegP initiates regulated processing of filamentous hemagglutinin in *Bordetella bronchiseptica*

**DOI:** 10.1101/2021.05.21.445233

**Authors:** Richard M. Johnson, Zachary M. Nash, Margaret R. Dedloff, John Shook, Peggy A. Cotter

**Author notes:** Address correspondence to Peggy A. Cotter.

## Abstract

Filamentous hemagglutinin (FhaB) is a critical virulence factor for both *Bordetella pertussis*, the causal agent of whooping cough, and the closely related species *Bordetella bronchiseptica*. FhaB is an adhesin, suppresses inflammatory cytokine production, and protects against phagocytic cell clearance during infection. Regulated degradation of the FhaB C-terminal prodomain is required to establish a persistent infection in mice. Two proteases, CtpA in the periplasm and SphB1 on the bacterial surface, are known to mediate FhaB processing, and we recently determined that CtpA functions before, and controls the FhaB cleavage site of, SphB1. However, the data indicate that another periplasmic protease must initiate degradation of the prodomain by removing a portion of the FhaB C terminus that inhibits CtpA-mediated degradation. Using a candidate approach, we identified DegP as the initiating protease. Deletion of *degP* or substitution of its predicted catalytic residue resulted in reduced creation of FHA′, the main product of FhaB processing, and an accumulation of full-length FhaB in whole cell lysates. Also, FHA′ was no longer released into culture supernatants in *degP* mutants. Alterations of the FhaB C terminus that relieve inhibition of CtpA abrogate the need for DegP, consistent with DegP functioning prior to CtpA in the processing pathway. DegP is not required for secretion of FhaB through FhaC or for adherence of the bacteria to host cells, indicating that DegP acts primarily as a protease and not a chaperone for FhaB in *B. bronchiseptica*. Our results highlight a role for HtrA family proteases in activation of virulence factors in pathogenic bacteria.

**Importance:** Two-partner secretion (TPS) systems are broadly distributed among Gram-negative bacteria and play important roles in bacterial pathogenesis. FhaB-FhaC is the prototypical member of the TPS family and here we identified the protease that initiates a processing cascade that controls FhaB function. Our results are significant because they provide insight into the molecular mechanism underlying the ability of *Bordetella* species to prevent clearance by phagocytic cells, which is critical for bacterial persistence in the lower respiratory tract. Our findings also highlight an underappreciated role for HtrA family proteases in processing specific bacterial virulence factors.

## Introduction

Gram-negative bacteria from the genus *Bordetella* cause highly contagious respiratory infections in mammals that have adverse impacts on public health and industrial agriculture(1). The human-specific pathogen *Bordetella pertussis* causes pertussis, also known as whooping cough, and is responsible for infecting an estimated 24 million children annually, causing 160,000 deaths(2). A majority of pertussis cases are from areas of the world with low (<85%) vaccine coverage. However, pertussis incidence has been increasing over the last 30 years even in areas where >90% of the population has received three doses of the dTap/Tdap vaccines, including the United States(3). The resurgence of pertussis correlates with the switch from whole-cell (wP) vaccines to acellular (aP) vaccines, which are composed of up to four purified *B. pertussis* proteins. While the increased incidence of pertussis is in part due to abiologic factors, such as improved diagnosis and case reporting, studies using baboons indicate that although wP and aP vaccines protect against disease, neither prevents *B. pertussis* colonization, and the aP vaccine also failed to prevent transmission(4).

The most widely used aP vaccine contains pertussis toxin, pertactin, fimbriae subunits, and a proteolytically processed version of filamentous hemagglutinin (FhaB) called FHA. FhaB is produced by all *Bordetella* species that infect mammals and is required for bacterial adherence to eukaryotic cells(5–7). Infection of small rodents with *Bordetella bronchiseptica*, a species closely related to *B. pertussis* that produces a nearly identical set of virulence factors, indicate that FhaB is functionally interchangeable between *B. bronchiseptica* and *B. pertussis* and that FhaB is required for persistent infection in the mammalian lower respiratory tract (LRT)(8–11). The mechanism by which FhaB mediates protection from inflammatory clearance is not understood(11).

FhaB is a member of the broadly distributed two-partner secretion (TPS, also known as Type Vb) family of bacterial secretion systems(12–14). It is synthesized as a 379 kDa preproprotein that possesses an extended N-terminal signal sequence that mediates Sec-dependent delivery to the periplasm(15, 16). Following removal of the signal sequence, the conserved N-terminal TPS domain of FhaB interacts with the periplasmic POTRA domains of its cognate transporter FhaC to begin translocation of FhaB across the outer membrane (OM)(17–19). While the FhaB N terminus remains anchored to FhaC, two-thirds of FhaB is secreted through FhaC in the N-to C-terminal direction, emerging on the bacterial surface as a hairpin consisting of a β-helical shaft that is topped by the globular mature C-terminal domain(20) (MCD) (Figure 1A). The remaining C-terminal third of FhaB (∼1,200 amino acids), called the prodomain (PD), is retained in the periplasm by the prodomain N terminus (PNT), which prevents further secretion through FhaC(20). FhaB undergoes complex regulated processing by multiple proteases. In *B. bronchiseptica* the tail-specific family member CtpA mediates degradation of the prodomain, resulting in the generation of the 250 kDa product FHA′, which stays membrane-associated temporarily but is ultimately released from the bacterial surface (Fig 1A). Under certain conditions (i.e., overnight growth in rich media) FHA′ can be cleaved by the surface-anchored exoprotease SphB1 to produce the 243 kDa product FHA (Fig 1A, dashed arrow), which is immediately released. The biological significance of SphB1-dependent cleavage is unclear. Although initially identified as a protease mediating release of FHA(21), it has subsequently been shown that FhaB polypeptides lacking the prodomain, such as FHA′, are efficiently released by both wild-type and SphB1-deficient strains of *B. bronchiseptica*(12, 22). The fully processed protein (i.e., FHA) was long considered the functional molecule, as the PD is thoroughly degraded and not detectable as a standalone protein(11). However, full-length FhaB plays a vital role during infection, as mutants lacking portions of the PD are rapidly cleared from the murine LRT, despite being able to adhere to host cells as effectively as wild-type *B. bronchiseptica*^11^. These findings suggest that full-length FhaB is required specifically for defense against inflammation-mediated clearance from the LRT, and we hypothesize that this activity requires properly regulated degradation of the FhaB PD.

**Figure 1.**
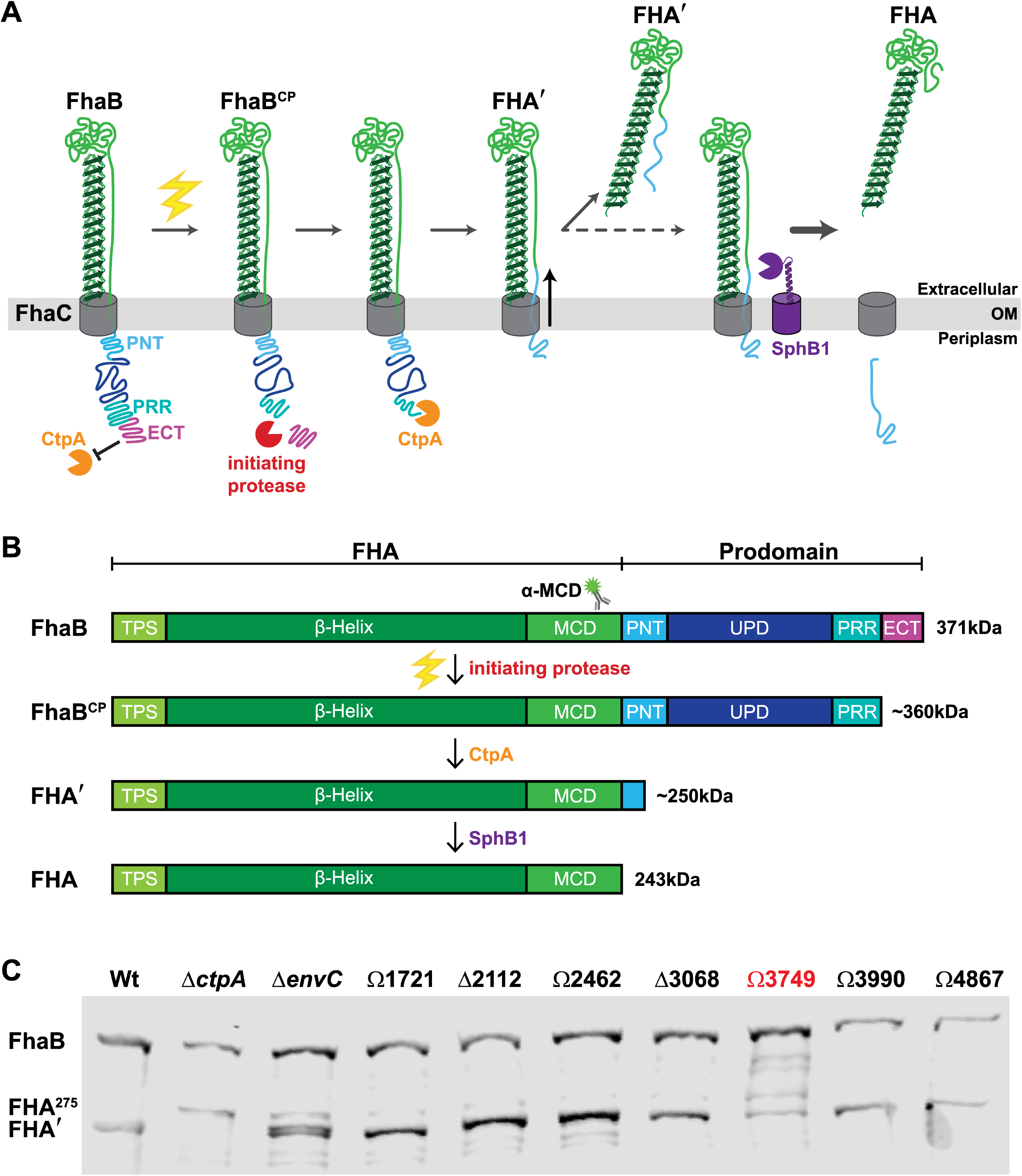
A screen of FhaB processing in *B. bronchiseptica* strains that carry mutations of predicted extra-cytoplasmic proteases identifies BB3749 as encoding a putative FhaB protease. (A) Our current model of regulated FhaB processing by three proteases. An unknown signal (electric bolt) causes the initiating protease to remove the Extreme C-terminus (ECT) that acts as a guard against CtpA. This clipped product (FhaB^CP^) is recognized by CtpA, which degrades most of the prodomain to form FHA’. Removal of a portion of the PNT by CtpA causes a shift in the molecule that allows the release of FHA’ into the supernatant. Under certain conditions this shift also exposes the primary SphB1 cleavage site and allows SphB1 to create FHA (dashed arrow), which is immediately released. (B) Linear schematics of FhaB (without the extended signal sequence) and the main polypeptides that result from processing by each protease. Molecular weights (kDa) are listed on the right. (C) Western blot analysis of whole cell lysates (WCL) from overnight cultures of *B. bronchiseptica* strains with deletions of (Δ) or plasmid disruptions in (Ω) the indicated ORFs. The antigenic region of the α-MCD polyclonal antibody is indicated by the cartoon of a fluorescent labeled antibody in (B).

Previous work from our laboratory determined that the last 98 amino acids of FhaB, called the ‘extreme C-terminus’ (ECT), negatively regulate CtpA-dependent FhaB processing(11). While characterizing CtpA, we identified a novel polypeptide intermediate called the “FhaB clipped product” (FhaB^CP^), which lacks ∼90 amino acids (i.e., most of the ECT) from the C terminus(22). FhaB^CP^ is stabilized in whole-cell lysates (WCLs) harvested from *B. bronchiseptica* Δ*ctpA* cultures, consistent with the discovery that the native FhaB C terminus is a poor substrate for CtpA(22) and the hypothesis that FhaB^CP^ is the preferred substrate for CtpA. The presence of FhaB^CP^ in Δ*ctpA* WCLs indicates that yet another protease, that we are calling the initiating protease, is responsible for converting FhaB into FhaB^CP^ through removal of the ECT. According to our model, CtpA degrades FhaB^CP^ to FHA′, which is either released from the bacterial surface or, if the bacteria are grown for extended periods of time, is cleaved by SphB1 to produce FHA, which is then immediately released (Figure 1A). We hypothesize that cleavage of the PD to form FhaB^CP^ is the critical control point as it initiates PD degradation. The goal of this study was to identify the initiating protease and to determine its contribution to FhaB processing and function.

## Results

### BB3749 (*degP*) appears to encode the initiating protease

To identify candidates for the initiating protease we searched the genome of wild-type *B. bronchiseptica* strain RB50 for ORFs predicted to encode proteases containing N-terminal Sec or TAT signal peptides. The 11 ORFs identified include those encoding both of the previously characterized proteases involved in processing FhaB, *sphB1*(21) and *ctpA*(22), as well as nine predicted to encode endopeptidases and carboxy-terminal acting peptidases (Table 1). We generated *B. bronchiseptica* strains containing either an in-frame deletion (ΔBB0299, ΔBB3068, ΔBB2112) or an integrated plasmid disruption (ΩBB1398, ΩBB1721, ΩBB2462, ΩBB3749, ΩBB3990, ΩBB4867) in one of the nine ORFs and examined FhaB/FHA’ in whole cell lysates (WCL) using α-MCD specific antisera (Figure 1C). We included the strain carrying an in-frame deletion of *ctpA* (Δ*ctpA*) that has been previously shown to be required for normal FhaB processing as a control (Figure 1C). Instead of processing FhaB to the 250kDa FHA′ product, the Δ*ctpA* strain converts FhaB to the larger FHA^275^ polypeptide, likely due to aberrant processing by as yet unidentified proteases(22). Two of the mutants displayed altered FhaB processing compared to the wild-type strain. BB0299 (*envC*) is located directly 5′ to, and is co-transcribed with, *ctpA*(22) and is predicted to encode an M23 family peptidase with a degenerate LytM (dLytM) domain. dLytM homologs in other species, such as EnvC in *E. coli*, serve as accessory factors that allosterically activate cell-wall cleaving hydrolases(24, 25). The phenotype of the Δ*envC*_*Bb*_ mutant is similar, but not identical, to that of the Δ*ctpA* strain, suggesting that EnvC_*Bb*_ may function as an accessory factor promoting CtpA activity (Figure 1C). As *envC* is in an operon with *ctpA*, it is also possible that the in-frame deletion mutation in *envC* could alter expression of *ctpA*. Characterizing the precise role of EnvC_*Bb*_ in FhaB processing is beyond the scope of the current work and will be addressed with future studies.

The strain containing a disruption in BB3749 (ΩBB3749) displayed the expected phenotype for loss of initiating protease activity; a majority of the FhaB protein remained as the full-length ∼371 kDa form (Figure 1C). We observed partially degraded FhaB polypeptides in ΩBB3749 WCLs that are not present in lysates of wild-type bacteria, suggesting that in the absence of the BB3749 gene product, unidentified proteases inefficiently degrade FhaB following overnight growth. BB3749 is predicted to encode a high-temperature requirement A (HtrA) family serine protease. Prokaryotic HtrA proteases are extra-cytoplasmic enzymes initially identified as factors necessary for bacterial survival at elevated temperatures(26–28). They contain trypsin-like serine protease domains at their N termini, and either one or two protein-binding PDZ domains at their C termini. The protein encoded by BB3749 is predicted to contain a serine protease domain with a conserved catalytic triad (a GNSGG motif and nearby histidine and aspartic acid residues), as well as two PDZ domains (Fig. 2A). Although annotated as a homolog of *mucD*, which encodes a protease responsible for regulating mucoidy (due to alginate overproduction) in *Pseudomonas aeruginosa* (Fig. S1), BB3749 is not associated with mucoidy in *B. bronchiseptica*. Moreover, the amino acid sequence of the protein encoded by BB3749 is 99.8% identical to that of *B. pertu*ssis DegP(29). Hence, we will hereafter refer to BB3749 as *degP* (or *degP*_*Bb*_ when distinguishing it from homologs is necessary) and its protein product as DegP (or DegP_*Bb*_).

**Figure 2.**
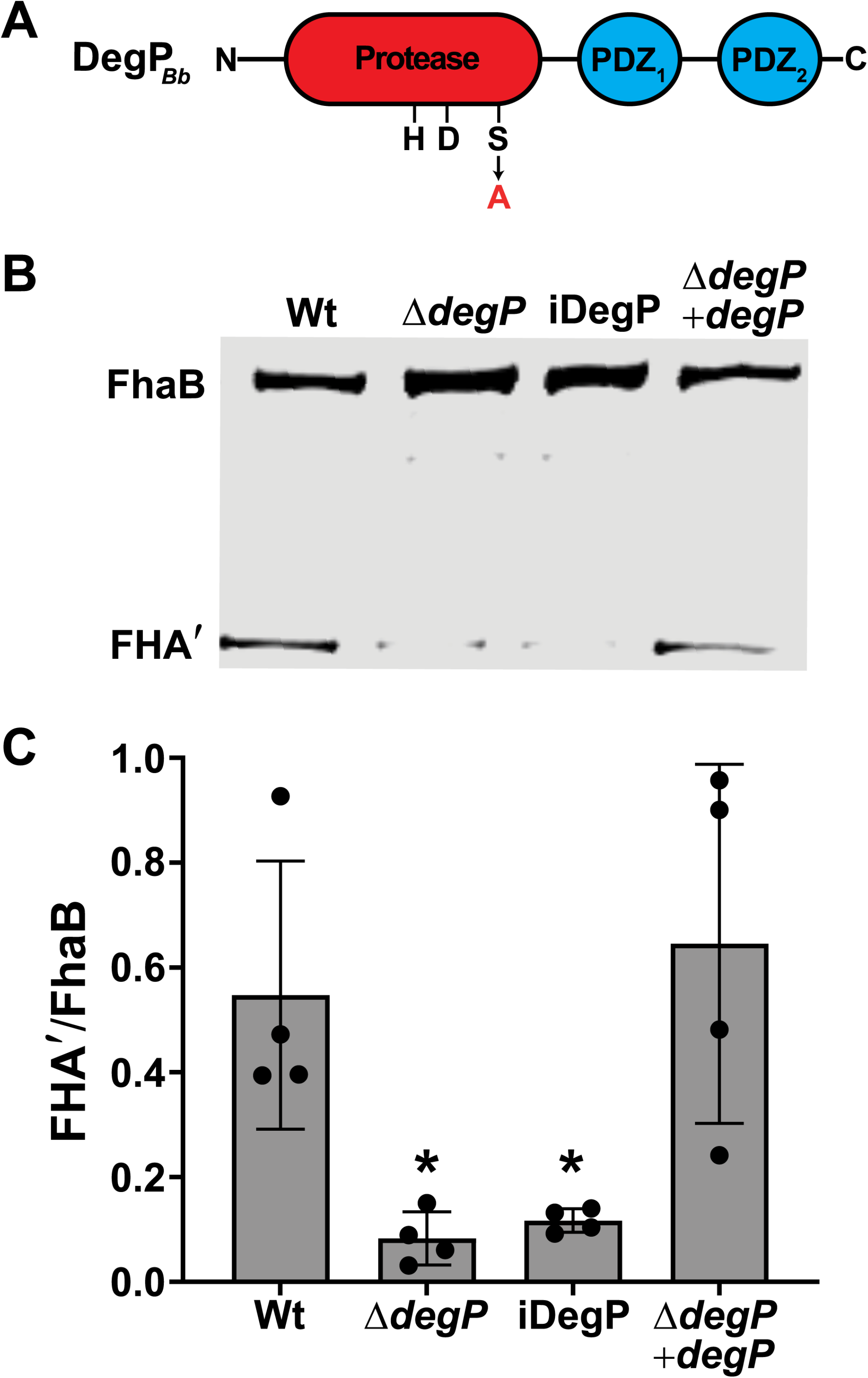
BB3749 encodes the HtrA family serine protease DegP, and its catalytic activity is required for efficient FhaB processing. (A) Diagram of DegP_*Bb*_ with the catalytic triad (HDS) and two PDZ domains indicated. The catalytic serine at position 237 was changed to an alanine to generate the inactive (iDegP) strain. (B) Western blot analysis of WCL from cultures grown for four hours in BvgAS-activating medium (SS) after overnight growth in BvgAS-inactivating medium (SS + 20mM MgSO_4_). (C) Ratio of processed FHA’ to full-length FhaB in the western blot in (B) and three similar blots. Significance of p<0.05 as determined by unpaired two-tailed T-test indicated by asterisk.

To determine if the FhaB processing defect displayed by the Ω*degP* strain was due to the loss of DegP (and not polar effects of the integrated plasmid), we constructed a strain in which codons 6-491 of *degP* were deleted (Δ*degP*). We also created a derivative of Δ*degP* in which a wild-type copy, driven by the constitutive S12 promoter, was inserted at the *att*Tn*7* site (Δ*degP* +*degP*). We grew the bacteria overnight in media containing 20mM MgSO_4_ (which inactivates the BvgAS regulatory system that activates expression of *fhaB* and other virulence factor-encoding genes) and then outgrew the bacteria in media without MgSO_4_ (BvgAS-activating conditions) for four hours to induce production of FhaB before collecting WCLs as previously described(22). As with the Ω*degP* strain, nearly all of the FhaB protein in the Δ*degP* strain was the full-length ∼371 kDa form, and normal processing was restored in the complemented strain (Fig. 2B and 2C). The intermediate FhaB products that were apparent in WCLs from the ΩBB3749 overnight cultures, were less prominent in the WCLs from Δ*degP* after four hours of growth under BvgAS-activating conditions, indicating that these aberrant products accumulate only after extended periods of growth (Fig 2B). For the remainder of this study we examined FhaB processing following four hours of growth under BvgAS-activating conditions.

To determine if the catalytic activity of DegP is required for FhaB processing, we constructed a strain (iDegP) in which codon 237 was changed to encode alanine instead of the predicted catalytic serine. The FhaB/FHA′ profile of the iDegP mutant was nearly identical to that of the Δ*degP* mutant, indicating that Ser237, and most likely the proteolytic activity, of DegP is required for efficient FhaB processing (Figure 2B and 2C).

### DegP is required for *B. bronchiseptica* growth at elevated temperatures

DegP homologs from other bacterial species, including *B. pertussis*(29), are required for bacterial survival and growth at elevated temperatures. To determine if *B. bronchiseptica* also requires DegP to grow under heat stress, we inoculated BG blood agar plates with wild-type, Δ*degP*, iDegP, and Δ*degP* +*degP* strains and incubated them at either 37°C or 42°C for 72 hours. Both the Δ*degP* and iDegP mutant strains were attenuated for growth at 42°C but not at 37°C (Fig. S2). Failure of the Δ*degP* strain to grow at 42°C was restored by expression of a wild-type copy of *degP in trans* (Δ*degP* +*degP*, Fig. S2). These data indicate that DegP, and its catalytic activity, are required for *B. bronchiseptica* to grow at 42°C. BvgAS activates (either directly or indirectly) the expression of all genes encoding *Bordetella*’s known protein virulence factors, including *fhaB*(30). Many of these factors are surface-localized or secreted into the extracellular environment and most, if not all, are produced at a high level when BvgAS is active. The severe growth defect displayed by *degP* mutants in response to elevated temperature was greatly alleviated in bacteria grown in the presence of 50 mM MgSO_4_, which inactivates BvgAS. Magnesium ions (Mg2+) are crucial for envelope stability(31), so we also grew all four strains on plates containing 50mM MgCl_2_. Unlike MgSO_4_, which rescued the growth of the mutant strains, addition of MgCl2 had no effect on bacterial growth at 42°C. These data suggest that the requirement for DegP under heat stress conditions is primarily due to its proteolytic activity on one or more BvgAS-activated virulence factors. (29)(32)

### DegP acts first in stepwise processing of the prodomain

Our current model for FhaB processing involves three proteases: two periplasmic enzymes, CtpA and the initiating protease, and one exoprotease, SphB1 (Figure 1A). We previously determined that CtpA and SphB1 function in series, with CtpA acting prior to, and determining the cleavage site of, SphB1(22). In fact, our data suggest that SphB1-dependent processing only occurs after extended time in culture and may have no functional relevance under normal (*in vivo*) circumstances. We have also shown previously that the ECT inhibits CtpA from degrading the PD, as deletion of the ECT (ΔECT) causes rapid and thorough conversion of FhaB to FHA′ (which we call hyperprocessing) in a CtpA-dependent manner(11, 22). We hypothesized that the initiating protease is responsible for removing the ECT to convert full-length FhaB into the CtpA substrate FhaB^CP^. To test this hypothesis, we first showed (as we have done previously(22)) that addition of an HA epitope N-terminal to the PRR (Fig. 3A, strain FhaB_HA-PRR_) has no effect on FhaB processing (Fig. 3B, both full-length FhaB and FHA′ are present in WCL of the FhaB_HA-PRR_ strain). Deletion of *degP* in this strain resulted in an accumulation of full-length FhaB_HA-PRR_ and a reduced amount of FHA′, just as it did in wild-type bacteria (Fig. 2), and complementation with wild-type *degP* restored FhaB processing (Fig. 3B). Deletion of *ctpA* does not alter the abundance of full-length FhaB in WCL, and results in the formation of FHA^275^ (Fig. 3B and(22)). (11, 22)FHA^275^ is a ∼275 kDa polypeptide containing an incompletely degraded PD. We next examined if *degP* is required for the hyperprocessing of FhaB that occurs in the absence of the ECT. In wild-type bacteria producing FhaB lacking the ECT (FhaB_ΔECT_, Fig. 3A), full-length FhaB is barely detectable (Fig. 3C). The same was true for the Δ*degP* strain (Fig. 3C), indicating that DegP does not contribute to the hyperprocessing of FhaB lacking the ECT. By contrast, FhaB_ΔECT_ was abundant in the Δ*ctpA* strain (Fig. 3C, yellow band as this strain also contains an HA epitope immediately N-terminal to the PRR), showing that the hyperprocessing of FhaB_ΔECT_ is CtpA-dependent. These data indicate that DegP acts prior to CtpA and is required for removal of the ECT to allow degradation of the prodomain by CtpA.

**Figure 3.**
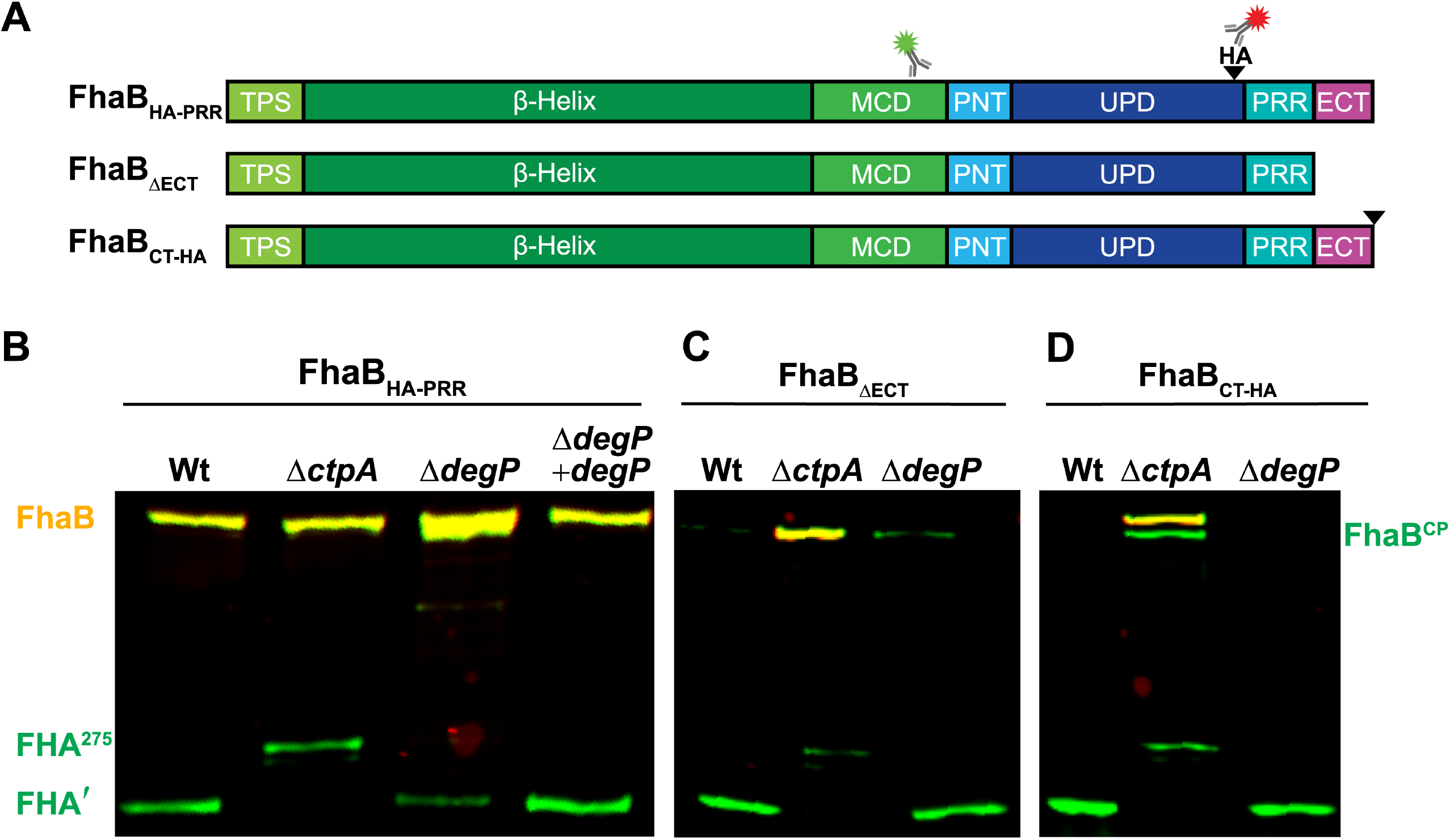
DegP removes the ECT to permit degradation of the prodomain by CtpA. (A) Linear schematic of FhaB_HA-PRR_, a version of FhaB that contains an HA tag (black wedge) inserted N-terminal to the proline rich region (PRR). The HA tag at this location is not disruptive to normal FhaB processing^1,2^. FhaB_ΔECT_ lacks the C-terminal 98 amino acids (ECT). FhaB_CT-HA_ has the HA tag located at the C terminus. (B) Western blot analysis of WCLs from strains grown for four hours in BvgAS-activating medium after overnight growth in BvgAS-inactivating medium. Full-length molecules (FhaB_HA-PRR_ and FhaB_CT-HA_ appear yellow due to recognition by both α-MCD (green) and α-HA (red) antibodies. The intermediate polypeptide FhaB^CP^ and final product FHA appear green due to loss of the HA tag. In the absence of CtpA, a 275 kDa polypeptide, FHA^275^, is created rather than FHA’.

Similar to deleting the ECT, and in contrast to inserting an HA epitope N-terminal to the PRR, adding an HA epitope to the C terminus of FhaB (FhaB_CT-HA_) results in CtpA-dependent hyperprocessing (Fig. 3D and(22)), i.e., this epitope somehow converts the FhaB C terminus into a substrate for CtpA. We reasoned that addition of the HA epitope to the C terminus could impact FhaB processing in two ways: 1) The HA epitope is itself could be a substrate for the tail-specific protease CtpA, and/or 2) Addition of the HA epitope could disrupt proper folding of the ECT causing it to be degraded by DegP. Deletion of *degP* did not prevent hyperprocessing of FhaB_CT-HA_, indicating that DegP is not required for CtpA-dependent degradation of the full-length FhaB_CT-HA_ and that CtpA proteolytic activity is not dependent on DegP. Consistent with CtpA-dependent degradation of FhaB_CT-HA_, we observed an accumulation of the full-length polypeptide (371 kDa yellow band) in WCLs from the Δ*ctpA* strain. We also detected the FhaB^CP^ in Δ*ctpA* WCLs, which we believe is the product formed following removal of the ECT by the initiating protease (∼362 kDa green band). The presence of the FhaB^CP^ in Δ*ctpA* FhaB_CT-HA_ and not in Δ*ctpA* FhaB_HA-PRR_ WCLs indicates that the addition of the C-terminal HA epitope promotes removal of the ECT-HA by the initiating protease (DegP). Taken together these data indicate that DegP acts first in stepwise processing of the PD.

### DegP is required for release of FhaB-derived polypeptides from the bacterial surface

*B. bronchiseptica* efficiently releases FhaB-derived polypeptides into culture supernatants and the specific polypeptide released depends on the age of the culture and the proteases produced by the bacterial strain (Fig. 4B). Wild-type *B. bronchiseptica* primarily releases FHA′ during the first 6 hours in culture under BvgAS-activating conditions, before SphB1 begins to cleave accumulated FHA′ to cause the formation and immediate release of FHA(22) (Supp. Fig 3). In the absence of CtpA, FHA′ is not produced, and instead FhaB is converted to the aberrant FHA^275^ product, which is retained on the cell surface. FHA^275^ cannot be released due to the presence of the complete PNT. However, FhaB or FHA^275^ can be cleaved by SphB1 promiscuously at non-preferred sites to form the smaller polypeptides FHA_1/2_, which are immediately released(20, 22)(Fig. 4A and 4B). None of the processed polypeptides (FHA′, FHA, or FHA_1/2_) were detected in significant amounts in the Δ*degP* mutant, demonstrating that DegP is critically important for release of FhaB-derived polypeptides.

**Figure 4.**
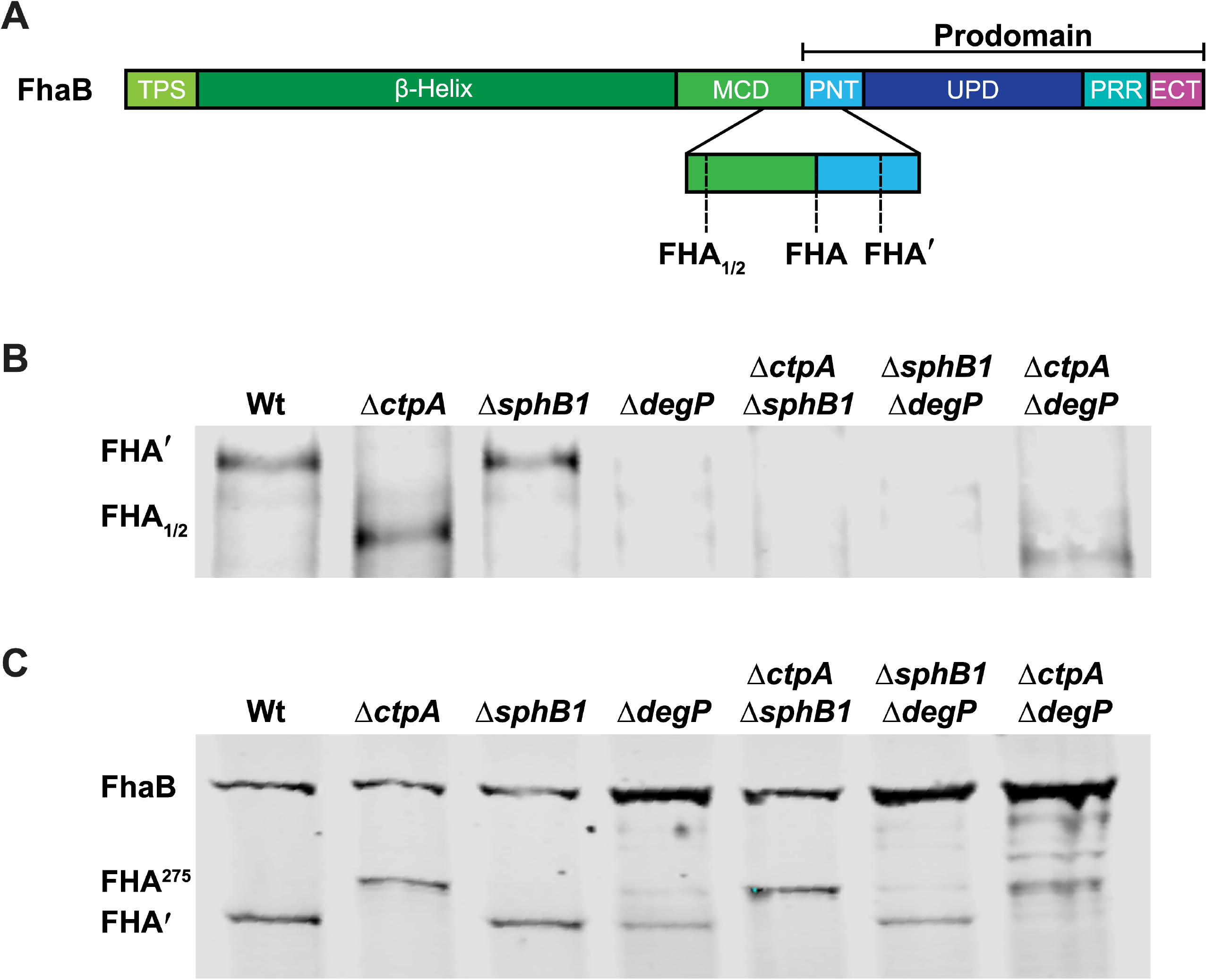
DegP_*Bb*_ is required for release of FhaB-derived polypeptides. (A) Diagram depicting the C termini of the FhaB-derived polypeptides released by *B. bronchiseptica*. Western blot analyses of (B) culture supernatants and (C) WCLs of strains grown for four hours in BvgAS-activating medium after overnight growth in BvgAS-inactivating medium. Blots probed with α-MCD antibody.

As shown in Fig. 4B and in our previous publication(22), CtpA <u>or</u> SphB1 is required for release of FhaB-derived polypeptides. In the absence of DegP, however, neither CtpA nor SphB1 is sufficient; FHA′ or FHA_1/2_ were barely detectable in supernatants of any strain containing the Δ*degP* mutation (Fig. 4B). These results reinforce the conclusion that DegP-mediated cleavage of FhaB occurs first and is required for subsequent degradation by CtpA and then cleavage by SphB1. When considered in conjunction with the WCL data (Fig. 4C), these data are also consistent with our model(12) (Fig. 1 and depicted in greater detail in Supplemental Fig 3.), which states that the polypeptide formed in the absence of CtpA (FHA^275^) cannot be released, but it can serve as a poor substrate for SphB1, cleavage by which generates FHA_1/2_ (because the PNT prevents movement through the FhaC channel that would allow the preferred SphB1 cleavage site to be exposed on the surface), which is immediately released (Fig. 4A and 4B). It is important to note that the supernatant samples have been concentrated substantially compared to the WCLs. Therefore, while levels of FHA_1/2_ in supernatants of the Δ*ctpA* strain appear to be similar to levels of FHA′ in supernatants of wild-type bacteria, no FHA_1/2_ was detected in WCLs of the Δ*ctpA* mutant and a substantial amount of FHA′ was present in WCLs of wild-type bacteria, indicating that in the absence of CtpA, generation of FHA_1/2_ by SphB1 is less efficient than generation of FHA′ by wild-type bacteria. In the absence of DegP, FHA^275^ is not formed and only a small amount of FHA′ is generated which cannot be detected in culture supernatants (Fig 4B).

### DegP is not required for translocation of FhaB through FhaC

DegP could be indirectly required for FhaB processing by promoting FhaB translocation through FhaC. To determine if loss of DegP prevents efficient translocation of FhaB to the cell surface, we used a dot blot assay to compare relative amounts of FhaB on the surface of intact bacteria following four hours of growth under BvgAS-activating conditions (Figure 5). In this assay, only surface-exposed epitopes are detectable by antibodies when cells are intact. When the bacteria are boiled to disrupt the membranes, internal epitopes can additionally be detected by antibodies. The α-MCD antibody (green) detected the MCD in unboiled bacteria, indicating that the MCD was located external to the outer membrane. However, the HA-tag was only detected after boiling the bacteria, indicating that the PD remains localized in the periplasm. As expected, neither the MCD nor the HA-tag was detected in the Δ*fhaB* strain. The protease mutants, Δ*ctpA* and Δ*degP*, were similar to wild-type bacteria in that the MCD was detected regardless of boiling while the HA-tag was only detected post-boiling. These results indicate that neither CtpA nor DegP is required for FhaB to be secreted through the outer membrane or for the PD to remain within the periplasm. Therefore, the reduction in FhaB processing that occurs in *degP* mutant strains is not due to mislocalization or instability of FhaB.

**Figure 5.**
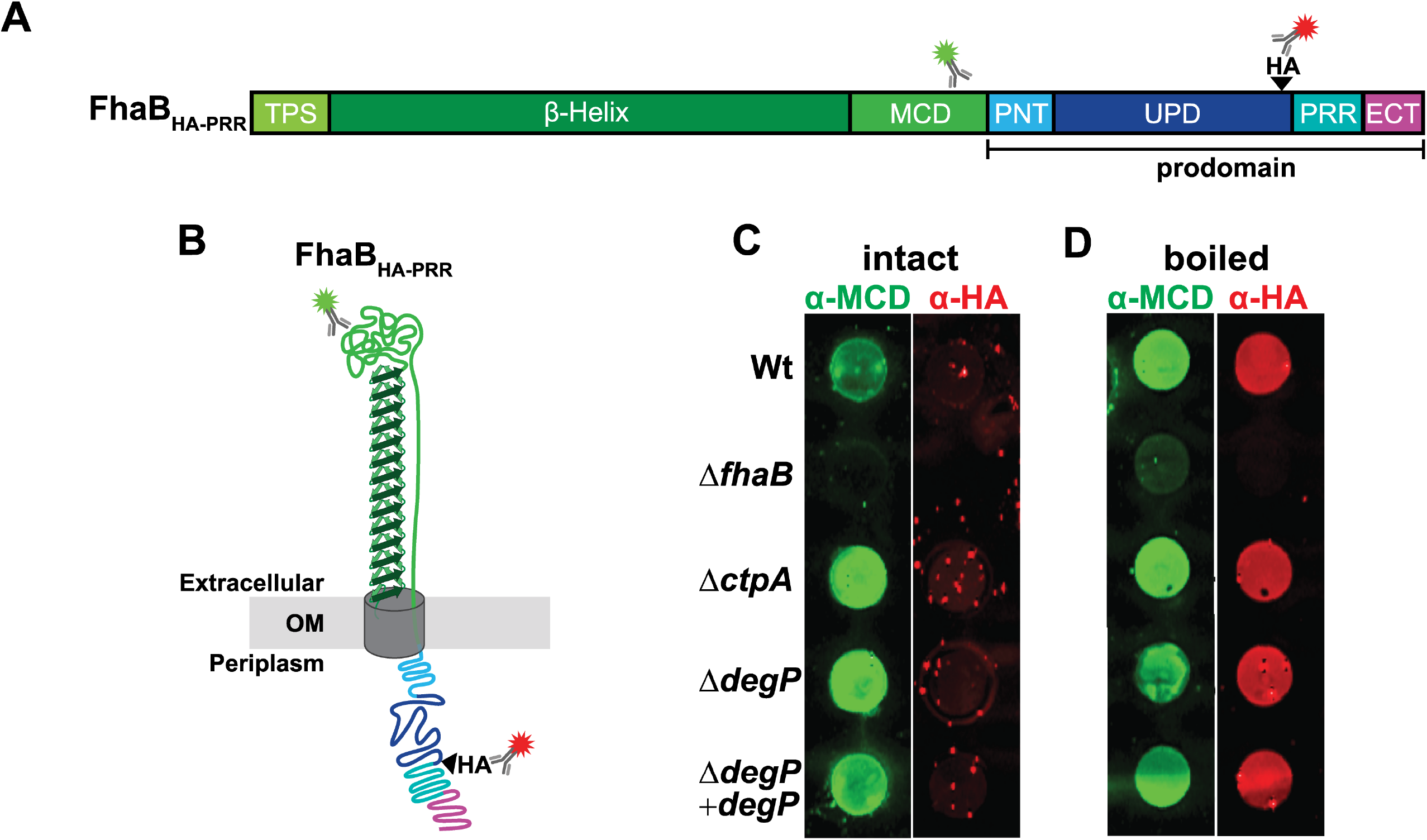
DegP is not required for FhaB secretion or retention of the prodomain within the periplasm. (A and B) Linear schematic and folded model of FhaB_HA-PRR_ with α-MCD and α-HA antibody binding regions shown. (C) Dot blots of intact and boiled cells. Blots were probed with α-MCD and α-HA antibodies.

### DegP is not required for FhaB-mediated adherence to host cells *in vitro*

While the dot blot analysis indicates that DegP is not required for translocation of FhaB to the bacterial surface, it cannot distinguish between active or inactive conformations of FhaB. We performed *in vitro* adherence assays to determine if the FhaB on the surface of Δ*degP* bacteria is capable of promoting bacterial adherence to human epithelial cells *in vitro*. Adherence of the Δ*degP* strain was similar to that of wild-type *B. bronchiseptica*, indicating that DegP is not required for translocation of FhaB to the bacterial surface or for the MCD to fold into an adherence-competent conformation (Figure 6). These data, along with the finding that strains that produce catalytically inactive DegP are defective for FhaB processing, are consistent with the primary function of DegP being that of a protease, rather than a chaperone, at least with regard to FhaB.

**Figure 6.**
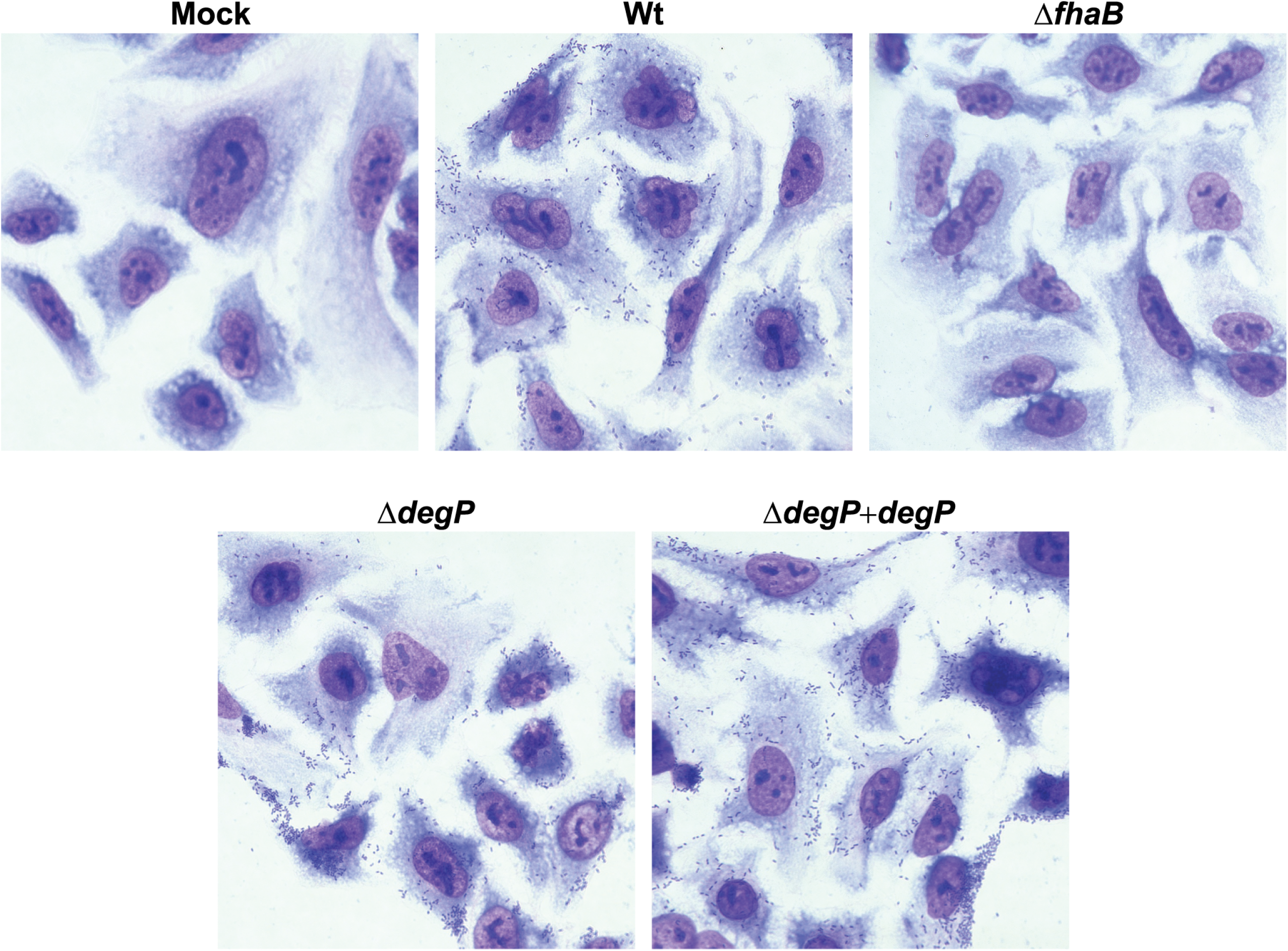
Deletion *of degP* does not disrupt FhaB-mediated adherence of *B. bronchiseptica* to epithelial cells *in vitro*. Representative images of Giemsa stained A549 human epithelial cells following a 15-minute inoculation with the indicated strains of *B. bronchiseptica* at an MOI of 100.

### DegP does not signal through the RseA-RpoE pathway to control FhaB processing indirectly

Some HtrA proteins, such as *E. coli* DegS (DegS_*Ec*_), are not degradative proteases. Instead, they transduce signals from the periplasm to cytoplasmic effectors, controlling the expression of genes encoding stress response factors indirectly. In the *E. coli* RseA-RpoE signal transduction system, the anti-sigma factor RseA, which is embedded in the inner membrane, binds and sequesters the RpoE sigma factor (σE) in the cytoplasm, preventing σE from binding to and directing RNA polymerase to activate expression of target genes(33). In *E. coli* undergoing exponential growth under standard laboratory conditions, DegS_*Ec*_ homotrimers associated with the inner membrane are in an inactive conformation. When *E. coli* is exposed to conditions that disrupt the folding and localization of proteins in the periplasm, misfolded proteins bind DegS_*Ec*_’s single PDZ domain, resulting in allosteric activation of DegS_*Ec*_’s proteolytic activity(34). When active, DegS_Ec_ cleaves a specific site on a periplasmic region of RseA, initiating a proteolytic cascade that causes the release of σE and the subsequent activation of stress response genes(33). Collectively this process is referred to as regulated intramembrane proteolysis (RIP)(35). Among the genes activated in the *E. coli* σE regulon is *degP*(36–39). Although the genomes of the *Bordetella* reference strains, Tohama I (*B. pertussis*) and RB50 (*B. bronchiseptica*), encode functional Rse-RpoE systems(40–42), they do not encode DegS homologs, as neither of the two HtrA homologs produced by these strains (DegP and the BB4867 gene product) are predicted to include TM domains capable of inserting in the inner membrane. Therefore, the protease responsible for initiating regulated proteolysis of RseA in *Bordetella* is not known. Unlike *E. coli*, in which *degP* is located distally on the chromosome from *rpoE, degP* is in an operon with *rpoE, rseA*, and *rseB* in *B. pertussis* and *B. bronchiseptica* (Figure 7A). These observations led us to ask if DegP could regulate FhaB processing indirectly, by cleaving RseA_*Bb*_ and subsequently increasing σE-dependent expression of a gene encoding yet another protease. RpoE is thought to be essential in *B. pertussis*(41) but is dispensable for growth of *B. bronchiseptica* under standard laboratory conditions(42). To determine if DegP signals through the RseA-RpoE pathway, we deleted *rpoE* in wild-type, Δ*degP*, and iDegP *B. bronchiseptica* (Figure 7B). If DegP signals through RseA-RpoE to activate expression of a gene encoding another protease, then loss of RpoE would cause the Δ*rpoE* strain to exhibit a similar defect in FhaB/FHA processing seen in WCLs from Δ*degP* and iDegP mutants. Instead, we observed that deletion of Δ*rpoE* had no effect on FhaB processing by any of the strains under the conditions examined, indicating that DegP is not indirectly regulating degradation of the FhaB PD via RpoE (Figure 7).

**Figure 7.**
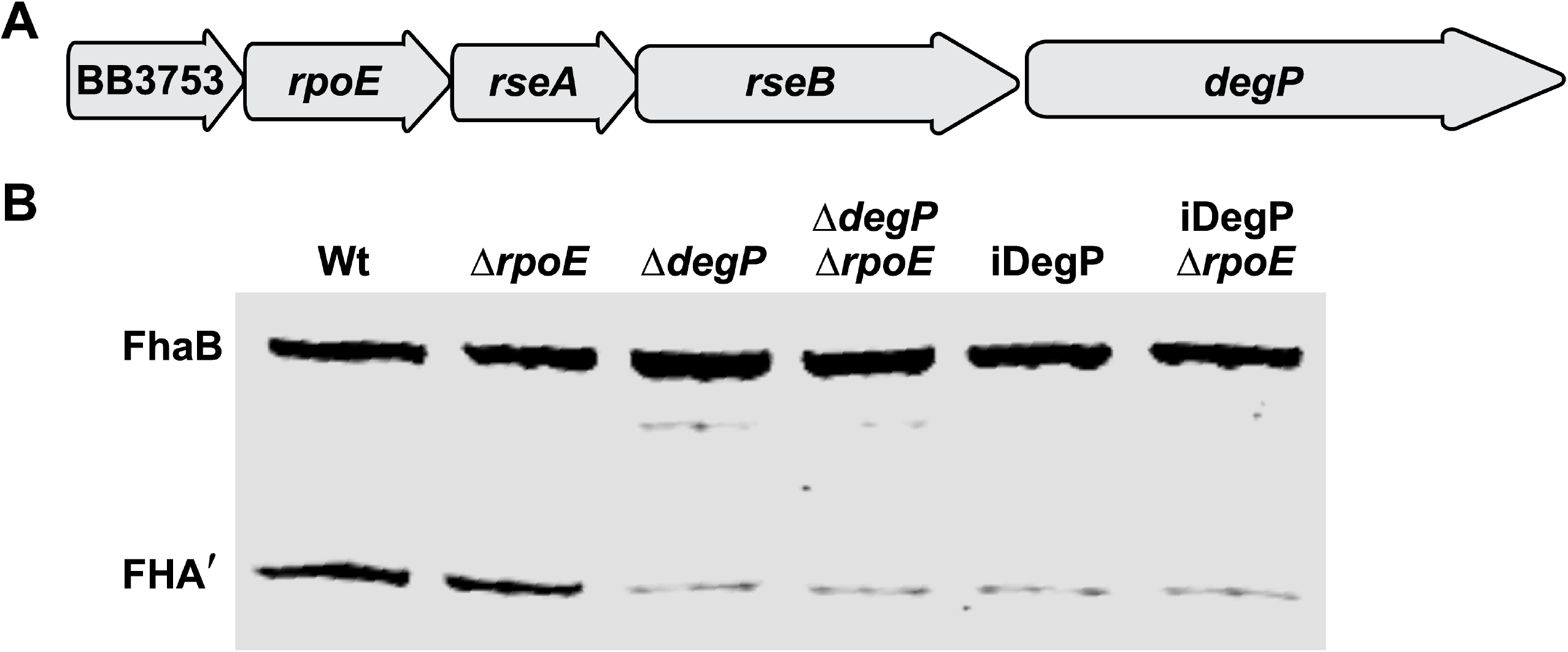
RpoE is not required for FhaB processing in *B. bronchiseptica*. (A) Diagram of the BB3753-BB3749 (*degP*) operon in the *B. bronchiseptica* RB50 genome. (B) Western blot analysis of WCL of strains grown for four hours in BvgAS-activating medium following overnight growth in BvgAS-inactivating medium. Blots probed with α-MCD antibody.

## Discussion

Our results indicate that the HtrA family protease DegP initiates degradation of the FhaB prodomain in *B. bronchiseptica*. Three key findings support this conclusion: 1) FhaB processing was dramatically reduced, but not completely abrogated, in strains lacking DegP (Δ*degP*) or producing DegP in which the predicted catalytic serine was replaced with alanine (S237A, iDegP). The small amount of FHA′ detected in WCLs of *degP* mutants was likely due to the prodomain being converted to a CtpA substrate by housekeeping proteases, which is revealed only in the absence of DegP. 2) DegP was dispensable for degradation of FhaB polypeptides with altered C termini (ΔECT and CT-HA) that are efficiently degraded by CtpA (Fig. 3). These data are consistent with our previous studies that determined that the native FhaB C terminus is a poor substrate for CtpA(22), and indicate that, in wild-type bacteria, DegP cleaves the prodomain first, generating a C terminus that is then efficiently degraded by CtpA. CtpA-dependent degradation of the ΔECT and CT-HA polypeptides in the absence of DegP also indicate that accumulation of full-length in WCLs from *degP* mutants is not due to the loss of CtpA activity. 3) DegP was not required for translocation of FhaB through FhaC (Fig. 5) or for FhaB to mediate adherence to epithelial cells (Fig. 6), consistent with DegP functioning as a protease and not a chaperone for FhaB. Previous studies determined that if FhaB is stalled in the periplasm (i.e., in *fhaC* mutants), it is degraded(43). The stability of full-length FhaB and the surface-localization of the MCD in *degP* mutants, therefore, indicate that DegP is not required to preserve FhaB stability or to deliver the TPS domain to FhaC in the periplasm. Although it is formally possible that DegP regulates FhaB processing indirectly by activating another protease, the most parsimonious explanation for our findings is that DegP acts on FhaB directly.

FhaB is required to mediate adherence to mammalian cells, prevent bacterial clearance by innate immune effectors, and control inflammatory signaling during infection. While processed FHA′ is sufficient for adherence, full-length FhaB and regulated degradation of the prodomain is required specifically for *B. bronchiseptica* to resist phagocytic cell clearance; mutants with altered prodomain processing are cleared rapidly from the lower respiratory tract but are not altered for adherence *in vitro* or *in vivo* or for suppression of inflammatory cytokines(11). Consistent with these data, the Δ*degP* strain adhered to human A549 epithelial cells as efficiently as wild-type bacteria do (Fig. 6), confirming that regulated prodomain processing is not required for FhaB to function as an adhesin. The fact that FhaB was exported to the cell surface (Fig. 5) and able to mediate adherence in *degP* mutants (Fig. 6) indicates that DegP does not function as a chaperone required to stabilize FhaB in the periplasm or to bring the N terminus of the TPS domain to FhaC. Furthermore, if DegP was an important chaperone for the FhaB prodomain, then lack of DegP would result in hyperprocessing, similar to the ΔECT strain. However, full-length FhaB was stabilized in *degP* mutants, indicating that DegP does not promote folding or stability of the prodomain.

Although *B. bronchiseptica* consistently produces and releases FHA′ (and sometimes FHA) from the bacterial surface during *in vitro* culture, neither the mechanism by which release occurs, nor its biological relevance, is understood. Our previous data suggested that either CtpA or SphB1 are sufficient for release(22). However, our new data indicate that their sufficiency is dependent on DegP as no FhaB-derived polypeptides are released by *degP* mutants (Fig. 4B). Future studies to determine how DegP and CtpA (and potentially EnvC) cooperate to degrade the prodomain should also reveal the mechanism underlying retention versus release of FhaB-derived polypeptides, another important outstanding question for TPS systems, in general.

DegP’s proteolytic activity is required for initiating FhaB processing in *B. bronchiseptica*, but it is not known whether the DegP homolog from *B. pertussis* is involved in degradation of the FhaB prodomain in that organism. A previous study that examined DegP in *B. pertussis* determined that this protease may have both catalytic and non-catalytic functions that are required for maintaining envelope integrity and are involved with FhaB secretion(29). However, that study did not examine prodomain degradation(29). Comparison between our study and Baud *et al*. highlights some key differences between *B. pertussis* and *B. bronchiseptica*, namely that *B. pertussis degP* mutants are more susceptible to elevated temperature than *B. bronchiseptica degP* mutants are. *B. pertussis degP* mutants are attenuated for growth at 37°C(29), while *B. bronchiseptica* strains lacking DegP present a growth defect at 42°C but not at 37°C (Supp Fig. 2).

Baud *et al*. determined that growth at 37°C of the *degP* mutants could be partially restored by expression of catalytically inactive DegP (S237A) *in trans*, which, combined with examination of binding interactions between DegP and non-native N-terminal FhaB polypeptides *in vitro*, led them to conclude that DegP functions as a chaperone in *B. pertussis*(29). By contrast, we found that DegP’s proteolytic activity is required for both FhaB processing and resistance to heat stress. Our iDegP strains, which produce DegP S237A from the native locus, were as attenuated for growth at 42°C and for FhaB processing as the Δ*degP* strain was (Supp Fig. 2 and Fig. 2), consistent with DegP in *B. bronchiseptica* functioning primarily as a protease for both FhaB processing and resistance to envelope stress.

In a follow up study, Baud *et al*. determined that DegP can be both soluble and associated with bacterial membranes as trimers, that membrane associated DegP has a higher affinity for non-native FHA polypeptides (lacking most of the MCD and all of the prodomain), and that membrane associated DegP has proteolytic activity(44). These findings provide insight into how DegP could be regulating degradation of the FhaB prodomain. We hypothesize that membrane associated DegP trimers could form a stable complex with the FhaB prodomain until a signal causes a conformational change in the prodomain such that DegP recognizes the prodomain as a substrate and initiates degradation. The signal that initiates degradation of the prodomain is unknown; however, previous work from our lab and others suggests that FhaB may be part of a novel toxin delivery system. The adenylyl cyclase toxin (ACT) – known to inactivate phagocytic cells(45–47) – binds FhaB on the bacterial surface(48, 49), and we hypothesize that regulated degradation of the FhaB prodomain controls delivery of surface-associated ACT to those cells. Our current model states that ACT forms a stable complex with full-length FhaB on the bacterial surface until these FhaB-ACT complexes interact with the ACT receptor (CR3, CD11b/CD18, Mac1) on phagocytic cells(46, 50). We propose that this interaction between FhaB-ACT and CR3 could generate a signal that is propagated through the FhaB MCD resulting in a conformational change in the prodomain allowing DegP to initiate degradation of the prodomain. Previous data indicates the FhaB ECT – which is 100% identical at the protein level between FhaB homologs in *B. pertussis, B. bronchiseptica* and *B. parapertussis –* likely adopts a conformation that stabilizes the prodomain by preventing degradation by CtpA(22). It is interesting to speculate that FhaB-ACT binding the CR3 receptor could trigger the ECT to unfold and therefore become a substrate for DegP, consistent with the preference for DegP homologs, such as those from *E. coli* and *B. pertussis*, to degrade unfolded proteins(29, 51). It is also possible that the signal that initiates FhaB processing is envelope stress that *B. bronchiseptica* faces during infection, which could induce conformational changes in the FhaB prodomain that cause it to become a substrate for DegP. Future studies will focus on identifying the signal that initiates FhaB processing and determining if regulated processing of FhaB is involved in ACT delivery to specific host cells.

## MATERIALS AND METHODS

### Bacterial strains

Bacterial strains and plasmids are listed in Supplemental Table 1. In-frame deletions were created by allelic exchange using derivatives of the pSS4245 plasmid(52). For complementation, *degP* was inserted into a derivative of the pUC18 plasmid that contained the S12 promoter and integrated into the chromosome at the *att*:Tn7 site(53). *E. coli* strains were used to amplify vectors (DH5α) and to transform *B. bronchiseptica* (RHO3). Mutations were confirmed by PCR and/or sequencing. As we have published in the past, our “wild-type” strain for FhaB studies is RBX11, a derivative of RB50 in which *fhaB* is more genetically tractable because the strain lacks *fhaS*, a gene highly homologous to *fhaB* but that plays no role in virulence(43).

### Culture media and conditions

*Bordetella bronchiseptica* strains were streaked from −80°C stocks onto Bordet-Gengou agar (BD Biosciences) supplemented with 6% defibrinated sheep blood (HemoStat Laboratories) and grown at 37°C for 2-3 days. Because some combinations of protease deletions, such as Δ*ctpA* Δ*degP*, resulted in growth defects that were minimized if BvgAS was not active, protease dual mutants were streaked onto blood plates supplemented with 50mM MgSO_4_. Colonies were picked from these plates and cultured overnight in Stainer-Scholte broth (SS) containing MgSO_4_ (50mM) to prevent production of Bvg-induced virulence factors, including FhaB. Bacteria were pelleted and washed with cold, sterile Dulbecco’s Phosphate Buffered Saline (DPBS, Gibco) and then cultured further for 4 hours in fresh Stainer-Scholte broth lacking MgSO_4_ to initiate production and processing of FhaB. This culture method is useful to examine early FhaB production and processing and to compare differences found across bacterial mutants that may be hidden after 16 hours of overnight growth(12). For experiments described in Figure 1, bacteria were simply grown overnight in Bvg inducing conditions (SS without MgSO_4_). *Escherichia coli* strains were grown at 37 °C in lysogeny broth or on lysogeny broth agar. Where appropriate, media was supplemented with streptomycin (20 μg/mL), kanamycin (50 μg/mL), and MgSO_4_ (50mM). For heat tolerance experiments (supplemental figure 2), the indicated strains were grown in Bvg+ phase liquid cultures (SS broth) overnight, normalized to 1 OD600 and then ten-fold serial dilutions of each culture were spotted (5uL per spot) onto BG blood agar plates with either 0 (Bvg+) or 50mM MgSO4 (Bvg–) and grown at 42 °C for 72 hours before being imaged.

A549 human epithelial cells were cultured as described by the supplier (ATCC) in Ham’s F12 media (Gibco) + 10% FBS (VWR). Cells were grown until they reached 80% confluency in T75 tissue culture treated flasks before passaging and the cells were passaged no more than 15 times out of a freeze and (Corning).

### Immunoblots

To examine cell-associated proteins by western blot, whole cell lysates were prepared by boiling pelleted cells in Laemmli buffer and shearing by passage through a 26G needle. For released proteins, culture supernatants were filtered through 0.2μm filters, and proteins were precipitated using 10% trichloroacetic, rinsed with cold acetone, resuspended in 1M Tris-HCl pH 8.8 and Laemmli buffer mixture, and boiled. Proteins were separated by SDS-PAGE using 4% or 5% polyacrylamide gels. Proteins were transferred to nitrocellulose membranes (GE Healthcare), and the membranes were probed with a rabbit polyclonal antibody generated against the FHA mature C-terminal domain (MCD)(10) and a mouse monoclonal antibody generated against an HA epitope (BioLegend). Corresponding α-rabbit and α-mouse IRDye secondary antibodies (LI-COR Biosciences) were used to detect proteins using a LI-COR Odyssey Classic Blot Imager (LI-COR Biosciences). Protein quantification was performed using LI-COR Image Studio version 5 software. *B. bronchiseptica* sample volumes were normalized based on optical density of the cultures.

To compare cell surface exoproteins versus internal proteins by dot blot, 100μL of 0.5OD_600_/mL of bacteria were spotted onto nitrocellulose membranes using a 96-well vacuum manifold. The membranes were probed using anti-MCD and anti-HA antibodies. Signal detected on intact bacteria corresponded to surface-exposed epitopes, and boiling disrupted the membrane to allow additional antibody interaction with otherwise inaccessible internal epitopes.

### Bacterial adherence assay

These assays were performed as previously described with minor alterations(5). A549 cells were seeded on sterile coverslips in 12-well plates. Once cells reached a confluency of 50% to 80%, the seeding medium was removed and the cells were inoculated with the indicated *B. bronchiseptica* strains (wt, Δ*fhaB, ΔdegP, ΔdegP* + *degP*) in SS broth at an MOI of 100 cfu per epithelial cell (2.5×10^7^ CFU/ml of bacteria). One set of wells were treated with SS broth alone (mock infection control). The bacterial density of each inocula were confirmed by plating on BG blood agar. The 12 well plates were centrifuged at 500 x g for 5 minutes at room temperature and then incubated for 15 minutes at 37 °C with 5% CO_2_. The inoculum was removed from each well and the cells were washed four times with Hanks’ Balanced Salt solution (Difco). Cells were then fixed with 0.5mL of ice-cold methanol for five minutes, after which the methanol was removed, and the coverslips were air dried for 10 minutes. The cells were then stained for 15 minutes with 0.5mL of 1:20 diluted Giemsa stain (Sigma Aldrich). The stain was removed, and coverslips were washed twice with water before being air dried and mounted onto slides with permount (Sigma Aldrich). The slides were imaged at 60X magnification (Keyence).

## Acknowledgements

We would like to thank the Cotter lab for helpful discussions, especially Jessica Beauchamp for critically reading the manuscript and providing editorial suggestions. We would like to thank the NIH for financially supporting this work, R01-AI129541, R01-AI153160, and the UNC Infectious disease and pathogenesis training grant T32-AI1007151. The funding agency had no role in study design, data collection and interpretation, or the decision to submit the work for publication. The authors report no conflicts of interest.

## Author contributions

RMJ, ZMN and PAC designed the studies, interpreted data and wrote the manuscript. RMJ, ZMN, MRD and JS acquired and analyzed the data.

---

**Figure S1. BB3749 encodes an HtrA family protease**.

(A) Alignment of the amino acid sequences of DegP of *B. bronchiseptica* (Uniport ID A0A0H3M188_BORBR), DegP of *B. pertussis* (Q7VW38_BORPE), MucD of *Pseudomonas aeruginosa* (G3XD20_PSEAE), DegP (DEGP_ECOLI), DegQ (DEGQ_ECOLI) and DegS (DEGS_ECOLI) of *E. coli*, and DegQ of *B. bronchiseptica* (AOAOH3LRV9_BORBR). Alignment was performed with Clustal Omega. (B) Matrix comparing the percent identities between the HtrA amino acid sequences

**Figure S2. DegP activity is required for *B. bronchiseptica* to grow at 42°C**. Ten-fold serial dilutions of *B. bronchiseptica* were spotted onto BG blood agar plates without magnesium additives (BvgAS-activating) or containing 50mM MgSO_4_ (BvgAS-inactivating) or 50mM MgCl_2_. Bacteria were grown for 72 hours at 37°C or 42°C as indicated.

**Figure S3. FhaB processing pathways of *B. bronchiseptica* protease mutants**. Linear diagrams of the model for regulated processing of FhaB in wild-type (A), Δ*ctpA* (B), Δ*degP* (C), Δ*sphB1* (D), Δ*ctpA*Δ*sphB1* (E), Δ*ctpA*Δ*degP* (F), and Δ*degP*Δ*sphB1* (G) strains. Polypeptides detected only in culture supernatants are indicated with a silcrow (§), polypeptides present in both WCL and supernatants are indicated with a psi (ψ), and polypeptides that are present in WCL in *degP* mutant strains, but at greatly reduced levels, are lightly shaded.

**Table S1.**
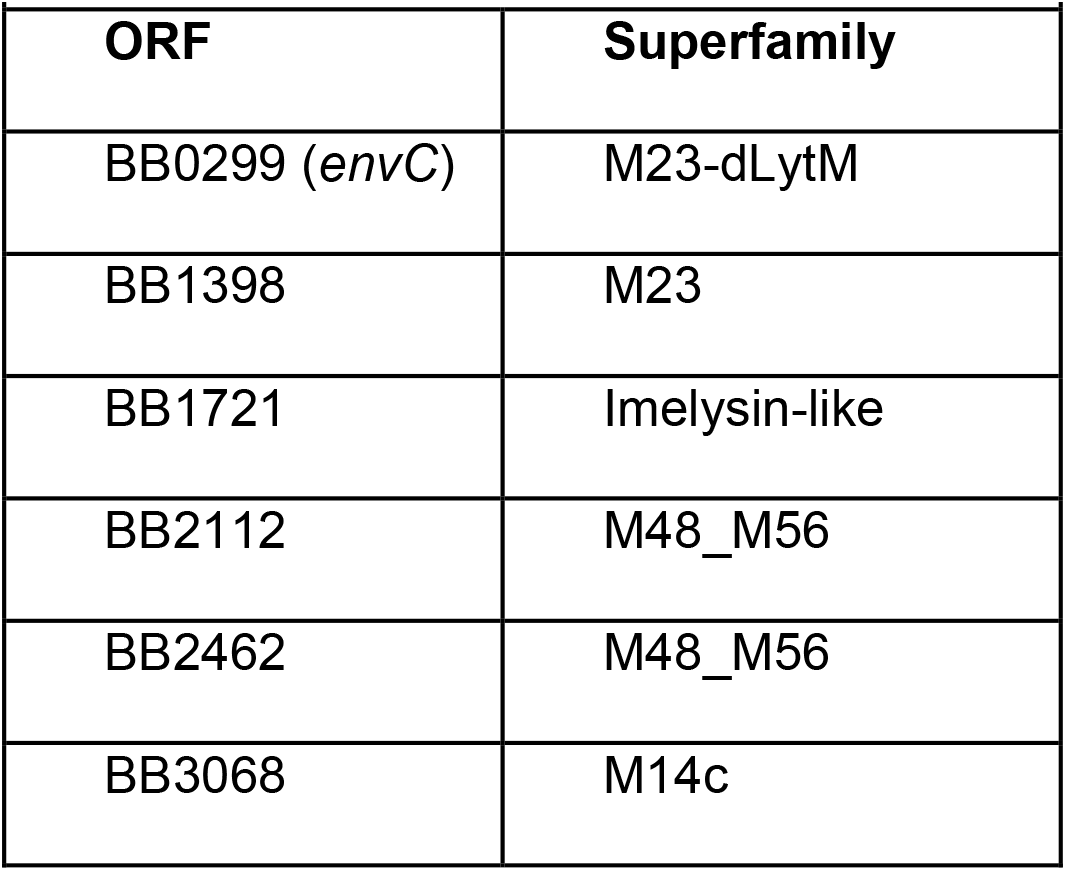
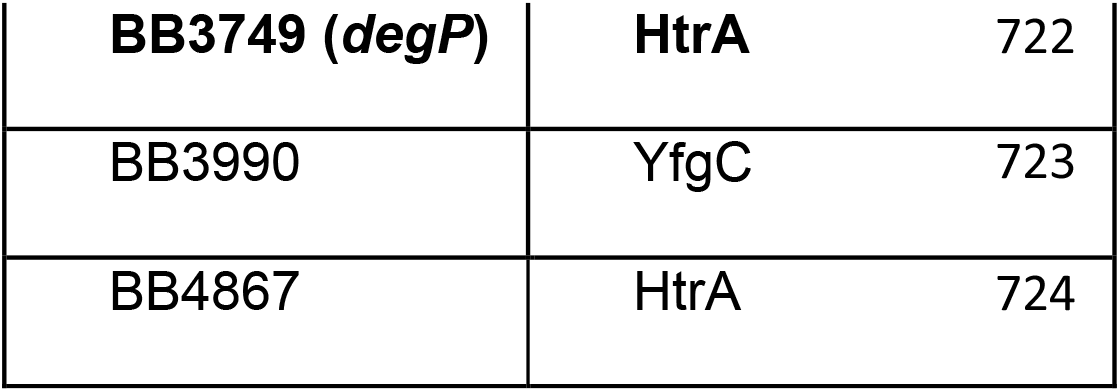
Strains and plasmids used during this study.

| Table 1. ORFs encoding initiating protease candidates |  |
| --- | --- |
| ORF | Superfamily |
| BB0299 (*envC*) | M23-dLytM |
| BB1398 | M23 |
| BB1721 | Imelysin-like |
| BB2112 | M48_M56 |
| BB2462 | M48_M56 |
| BB3068 | M14c |

|  |  |  |
| --- | --- | --- |
| **BB3749 (*degP*)** | **HtrA** | 722 |
| BB3990 | YfgC | 723 |
| BB4867 | HtrA | 724 |

